# *MicrobComp* – graphical user interface enables an *in-silico* comparison of taxonomic differences between two amplicon datasets

**DOI:** 10.1101/717231

**Authors:** Thomas Nussbaumer

**Affiliations:** Independent Researcher. Austria

## Abstract

The experimental and computational analysis of microbes is in the focus of basic and applied research for several decades already and has massively extended our knowledge of role and impact of microbes during that time.

Amplicon sequencing is still widely adapted and standard procedure to determine key species in a biological sample and despite a high number of different datasets being steadily generated and fed into current data archives, there is a shortcoming in data interpretation. Furthermore, more computational tools are still required to guide the researcher towards an in-depth understanding of an experiment in a more systematic way. Towards that aim, *MicrobComp* was created as a graphical user interface which allows to compare two separate species frequency tables that contain the relative or absolute abundance of microbial species in an experiment. Therefore, this tool can be of great help when studying different amplicon experiments where studies were originally performed in different experimental labs.

## INTRODUCTION

Microbes can be cause for life-threatening diseases (*e.g. Neisseria meningitidis* that can cause sepsis and meningitis [1] or Listeriosis which is caused by *Listeria monocytogenes* [2] among many more diseases where microbes play a central role. On the other side, microbes can have beneficial impact on the host when they counteract detrimental species’ by taking away necessary nutrients, by triggering or activating the immune system of the host [3-5] and can represent promising sources for future treatments for certain diseases. In addition, microbes are often dependent on the environmental milieu where they are supposingly harmless in certain tissues but harmful in another tissues [6].

Microbiome research has helped the scientific community to obtain a better functional insight into the coexistence with the host where the abundance of these species or their gene products can explain drastic gene expression profile changes in eukaryotic host genes [7]. This is possible by a bouquet of different experimental techniques such as dual transcriptomics, metatranscriptomics or metagenomics among other methods of the meta’omics collection [8]. These sequencing techniques have altogether or in various combinations helped to enable a more systemic or ‘holobiont view’ of all members that interact.

However, when it comes to the classification of a microbial species, amplicon sequencing is still the most accurate and most frequently applied way [9] but the annotation of species can be challenging: This is a consequence of the taxonomic annotation that relies on a small genomic fragment of a very conserved gene, the 16S rRNA gene, which includes nine hypervariable regions.

The selection of the flanking primer pairs that surround one or two hypervariable sites has also a big impact on the taxonomic resolution between different microbial genera. Fragments within the 16S rRNA gene can appear identical over several hundred base pairs even within genera. Still, nucleotide divergence might be hidden at other the remaining parts of the gene. This explains, why results are often different when different primer pairs and sequencing kits [10] are used and therefore a fair comparison of different techniques would require an almost identical experimental setups. This scientific challenge becomes more challenging as certain strains differ in their 16S rRNA copy numbers as well and low coverage datasets might not allow to resolve them especially when there is hardly any nucleotide divergence.

The annotations improved drastically with the availability of novel sequenced strains and release of many amplicon sequencing studies. Resources such as SILVA Arb [11] or eggNOG [12] are likewise needed to compare different study outcomes from amplicon sequencing projects. Furthermore, databases and projects which collect whole genome repositories such as NCBI Microbial Genomes [13]. If these resources are available, they also need to be consistently maintained.

One of the main challenges in today’s’ amplicon sequencing initiatives is the comparison of different studies that describe the same effect as in the current study. Typically, the main conclusions from the study are used for comparisons in favor of a remapping or reassembly of published data. These differences are also caused by differences in the quality of functional annotations which has benefited from the massive increase of new data over the last years. In addition, as most studies don’t contain both, the sequencing and frequency tables, this also hampers direct comparisons. There are many ways to compare OTUs tables but two of them are provided in the current tool.

Here, we demonstrate *MicroComp* which allows in a graphical interface to include two OTU tables and then compares them based on the existing functional annotation.

## MATERIAL AND METHODS

### Graphical user interface

The graphical user interface was implemented with help of the scripting language R and various scripts that are available under https://github.com/nthomasCUBE/microbComp. These scripts can be run with help of the R Shiny framework allowing an interactive comparison.

### Search for species of interest

A user can always compare two OTU tables where a comparison is performed by summing up or by the taking the mean (arithmetic mean or median) over all reads for one or many condition from one OTU table when compared to the OTU table and by selecting one taxonomic unit (e.g. family level or genus level).

## RESULTS

*MicroComp* provides an interface to integrate up to two microbial datasets that need to be available in form of a text file that includes the conditions as column names and OTUs as row names while the information of the taxonomic assignment is given in the last column and unique identifier of the OTU in the first column. The taxonomic assignment must include all taxonomic information from ‘kingdom’ level to ‘species’ level and needs to be separated by a semicolon. We used the data outcome from the study [14]describing microbiome data of IL-22 treated mice to demonstrate various features of the tool. **Figure 1** shows a comparison of two OTUs tables where we compared two samples based on the raw reads while also the relative frequency can be used and comparisons can be made based on all taxonomic levels, if the data is provided by the user. Furthermore, it is possible to compare groups of conditions from one OTU table versus groups of conditions from another OTU table (**Figure 2**). To support in this comparison, the sum, the average or the median can be taken along all conditions from the various samples groups. At last, for each group comparison a statistical test can be calculated by considering the relative frequencies along all conditions and by using a Wilcoxon test with or without adjustment for multiple testing (**Figure 3**).

**Figure 1.**
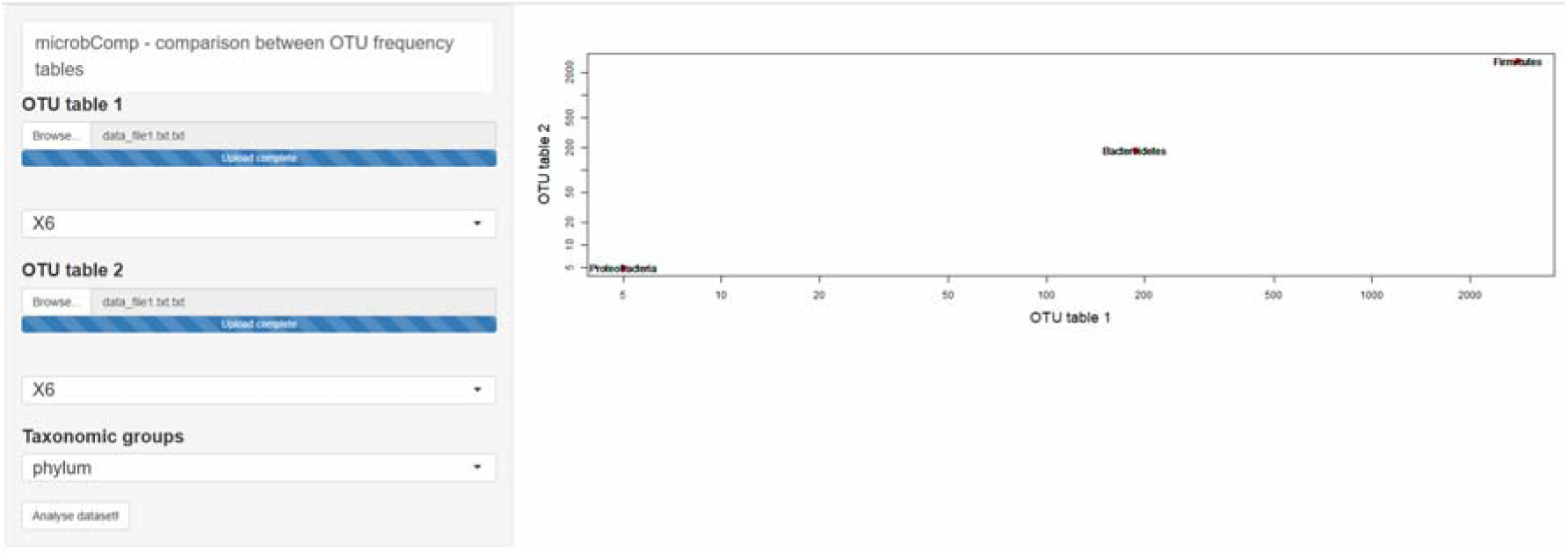
*MicrobComp* – graphical user interface with possibilities of the microbial datasets to include, analyses and visualization methods on phylum level.

**Figure 2.**
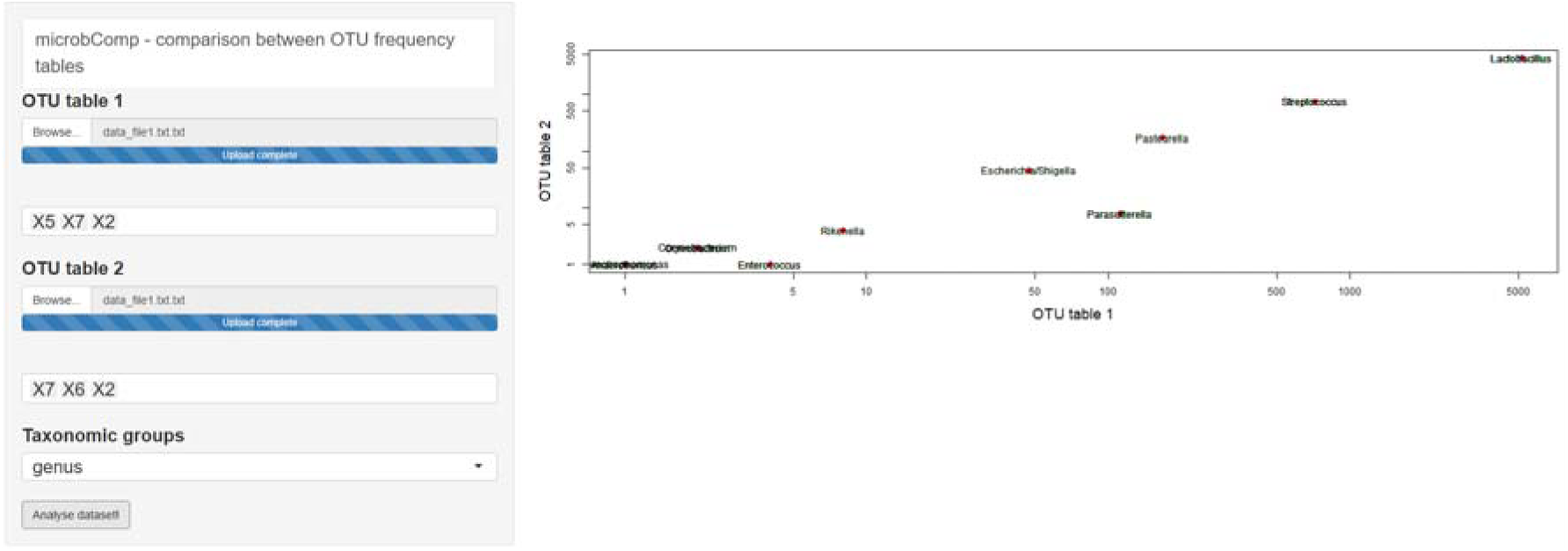
MicrobComp -graphical user interface with possibilities to compare groups of conditions versus other groups of conditions on genus level.

**Figure 3.**
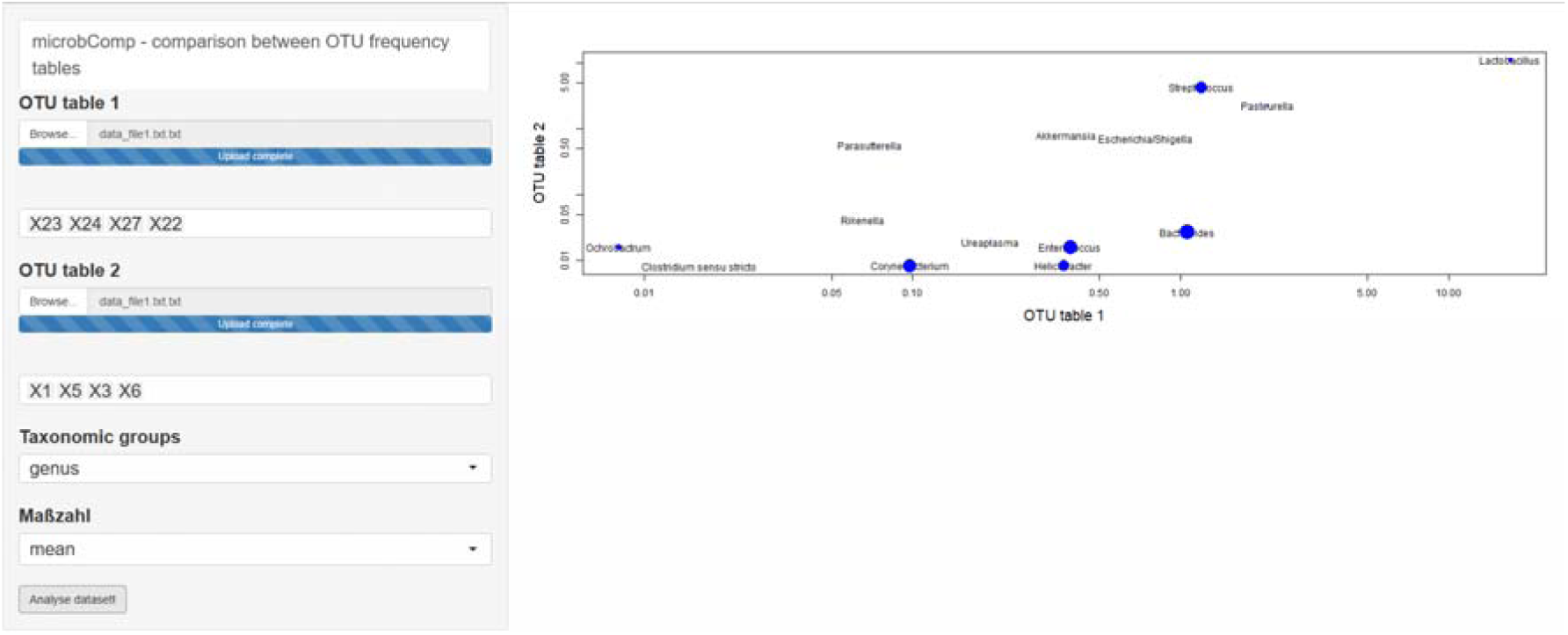
Statistical tests to determine differences in the relative frequencies of samples-groups between two different OTU frequency tables. Larger dots represent those where the p-value is closer 0 zero while no differences are observed when the dots are no shown.

## DISCUSSION

In order to compare different OTU tables efficiently it requires often methods that allow to determine commonalities or differences between a current study when compared to other studies that describe a similar experiment. Otherwise, such a comparison could be needed between different sequencing runs within a project, where researchers would like to understand whether species of interest are supported also by a more deeper sequenced run. To allow that in an efficient way, we generated *microbComp* to efficiently compare two OTU tables.

## REFERENCES

1. Stephens, David S., Brian Greenwood, and Petter Brandtzaeg. “Epidemic meningitis, meningococcaemia, and Neisseria meningitidis.” The Lancet 369.9580 (2007): 2196–2210.

2. Farber, J. M., and P. I. Peterkin. “Listeria monocytogenes, a food-borne pathogen.” Microbiology and Molecular Biology Reviews 55.3 (1991): 476–511.

3. Sassone-Corsi, Martina, and Manuela Raffatellu. “No vacancy: how beneficial microbes cooperate with immunity to provide colonization resistance to pathogens.” The Journal of Immunology 194.9 (2015): 4081–4087.

4. Pineda, Ana, et al. “Helping plants to deal with insects: the role of beneficial soilborne microbes.” Trends in plant science 15.9 (2010): 507–514.

5. Pii, Youry, et al. “Microbial interactions in the rhizosphere: beneficial influences of plant growth-promoting rhizobacteria on nutrient acquisition process. A review.” Biology and Fertility of Soils 51.4 (2015): 403–415.

6. Maggini, Valentina, et al. “Tissue specificity and differential effects on in vitro plant growth of single bacterial endophytes isolated from the roots, leaves and rhizospheric soil of Echinacea purpurea.” BMC plant biology 19.1 (2019): 284.

7. Huang, Yong, et al. “Microbes are associated with host innate immune response in idiopathic pulmonary fibrosis.” American journal of respiratory and critical care medicine 196.2 (2017): 208–219.

8. Segata, Nicola, et al. “Computational meta’omics for microbial community studies.” Molecular systems biology 9.1 (2013).

9. Hodkinson, Brendan P., and Elizabeth A. Grice. “Next-generation sequencing: a review of technologies and tools for wound microbiome research.” Advances in wound care 4.1 (2015): 50–58.

10. Guo, Feng, et al. “Taxonomic precision of different hypervariable regions of 16S rRNA gene and annotation methods for functional bacterial groups in biological wastewater treatment.” PloS one 8.10 (2013): e76185.

11. Quast, Christian, et al. “The SILVA ribosomal RNA gene database project: improved data processing and web-based tools.” Nucleic acids research 41.D1 (2012): D590–D596.

12. Huerta-Cepas, Jaime, et al. “eggNOG 4.5: a hierarchical orthology framework with improved functional annotations for eukaryotic, prokaryotic and viral sequences.” Nucleic acids research 44.D1 (2015): D286–D293.

13. Tatusova, Tatiana, et al. “NCBI prokaryotic genome annotation pipeline.” Nucleic acids research 44.14 (2016): 6614–6624.

14. Hernández, Pedro P., et al. “Interferon-λ and interleukin 22 act synergistically for the induction of interferon-stimulated genes and control of rotavirus infection.” Nature immunology 16.7 (2015): 698.

